# GM-CSF engages multiple signaling pathways to enhance pro-inflammatory cytokine responses in human monocytes during *Legionella* infection

**DOI:** 10.1101/2024.12.05.627084

**Authors:** Víctor R. Vázquez Marrero, Madison Dresler, Mikel D. Haggadone, Allyson Lu, Sunny Shin

## Abstract

The proinflammatory cytokine granulocyte-macrophage colony-stimulating factor (GM-CSF) is required for host defense against a wide range of pathogens. We previously found that GM-CSF enhances inflammatory cytokine production in murine monocytes and is required for *in vivo* control of the intracellular bacterial pathogen *Legionella pneumophila*. It is unclear whether GM-CSF similarly augments cytokine production in human monocytes during bacterial infection.

Here, we find that GM-CSF enhances inflammatory cytokine expression in *Legionella-*infected human monocytes by engaging multiple signaling pathways. *Legionella*- and TLR-dependent NF-𝜅B signaling is a prerequisite signal for GM-CSF to promote cytokine expression. Then, GM-CSF-driven JAK2/STAT5 signaling is required to augment cytokine expression in *Legionella*-infected human monocytes. We also found a role for PI-3K/Akt/mTORC1 signaling in GM-CSF-dependent upregulation of cytokine expression. Finally, glycolysis and amino acid metabolism are also critical for GM-CSF to boost cytokine gene expression in human monocytes during infection. Our findings show that GM-CSF-mediated enhancement of cytokine expression in infected human monocytes is regulated by multiple signaling pathways, thereby allowing the host to fine tune antibacterial immunity.

## Introduction

Respiratory infections are a leading cause of morbidity and mortality worldwide, as evidenced by the COVID-19 and tuberculosis pandemics (1, 2). More than 4 million people die annually due to respiratory infections (3). Continuous exposure to airborne pathogens has caused the respiratory mucosa to evolve sophisticated strategies to combat infections (4, 5). The lungs rely on innate immune cells and the airway epithelium to rapidly respond to disease-causing pathogens and secrete cytokines that orchestrate antimicrobial responses to control infection. One such cytokine known to be critical for control of lung pathogens is granulocyte-macrophage colony-stimulating factor (GM-CSF) (6).

GM-CSF was initially described as a growth factor due to its ability to differentiate hematopoietic precursors *in vitro* into myeloid cell populations, such as dendritic cells (DCs) (7). However, besides alveolar macrophages and some DC subsets that require GM-CSF for their development, GM-CSF does not otherwise appear to play a major role in steady state myelopoiesis *in vivo* (6, 8–11). It is now appreciated that GM-CSF is highly induced in a variety of diseases (6, 12) and can act as an inflammatory cytokine required to control several lung pathogens in mice, such as *Mycobacterium tuberculosis*, *Aspergillus fumigatus*, and influenza virus (13–17). Furthermore, human patients that develop autoantibodies against GM-CSF have increased susceptibility to infection (18). Thus, GM-CSF is an important driver of inflammatory responses and host defense. However, the mechanisms underlying how GM-CSF promotes inflammation during infection remains poorly understood.

One bacterial pathogen that is controlled by GM-CSF-driven immunity is *Legionella pneumophila*, a gram-negative bacterium that causes the severe pneumonia Legionnaires’ disease (19, 20). Upon entry into the lungs, *Legionella* infects and replicates within alveolar macrophages. *Legionella* uses a type IV secretion system that delivers effector proteins into the host cell cytosol to facilitate its intracellular survival and replication (21–24). A subset of these effectors potently inhibit host protein synthesis, thereby disabling the ability of macrophages to produce pro-inflammatory cytokines such as TNF and IL-12 (25–31). Despite this block in host translation, macrophages can still translate and release IL-1𝛼 and IL-1𝛽 (32, 33). We have shown that IL-1𝛼 and IL-1𝛽 are critical for control of *L. pneumophila* infection in mice (34, 35), in part by initiating a cellular communication circuit via GM-CSF (36). IL-1 receptor signaling in type II alveolar epithelial cells drives production of GM-CSF. Subsequently, cell-intrinsic GM-CSF receptor signaling in monocytes and other myeloid cells enabled their production of inflammatory cytokines during *Legionella* infection (36). Thus, GM-CSF promotes cytokine production in monocytes and other myeloid cells to orchestrate immune-mediated control of pulmonary *Legionella* infection in mice.

To investigate how GM-CSF promotes cytokine production in monocytes, we previously used primary murine monocytes infected with *Legionella ex vivo* as a model. We found that GM-CSF engages janus kinase 2 (JAK2)/signal transducer and activator of transcription 5 (STAT5) signaling to enhance aerobic glycolysis, which was required for maximal cytokine production in *Legionella*-infected monocytes (36). It is unclear whether a similar GM-CSF-driven pathway also enhances cytokine production in human monocytes. Most of the functional and mechanistic studies on the effects of GM-CSF in human monocytes have been conducted in the context of stimulation with purified lipopolysaccharide (LPS) (37–42). However, during bacterial infection, monocytes encounter a plethora of other bacterial-derived pathogen-associated molecular patterns (PAMPs) and signals in addition to LPS that may influence their function. Furthermore, *Legionella* and other bacterial pathogens employ virulence factors that interfere with protein translation and innate immune signaling that would be expected to limit cytokine production in monocytes. It is therefore important to study the effect of GM-CSF on human monocytes in the context of active bacterial infection.

In this study, we sought to mechanistically investigate the effect of GM-CSF on pro-inflammatory cytokine production in *Legionella-*infected human monocytes. We show that in both the THP-1 monocytic cell line and primary human monocytes, GM-CSF enhances *Legionella*-induced expression of the cytokines IL-1𝛼, IL-1𝛽, and IL-6. We find that *Legionella*-induced nuclear factor kappa B (NF-κB) signaling is a prerequisite signal for GM-CSF to enhance cytokine production in monocytes. We show that GM-CSF acts through JAK2-STAT5 signaling to enhance cytokine expression. Additionally, we find that phosphatidylinositol-3-kinase (PI-3K) signaling is required for GM-CSF to enhance *Legionella*-induced cytokine responses. Finally, both glycolysis and amino acid metabolism are required for GM-CSF to promote cytokine production in infected human monocytes. Altogether, these data demonstrate that GM-CSF-driven cytokine production is controlled by multiple signaling and metabolic pathways, thereby allowing the host to fine tune antibacterial immunity.

## Results

### GM-CSF enhances inflammatory cytokine production in human monocytes during

### Legionella infection

We previously found that GM-CSF enhances inflammatory cytokine expression in murine monocytes infected with *Legionella* (36). To determine whether GM-CSF also enhances cytokine production in human monocytes, we pretreated the human monocytic THP-1 cell line with recombinant GM-CSF prior to infection with *Legionella* or stimulation with the Toll-like receptor 2 (TLR2) agonist Pam3CSK4. We found that GM-CSF pretreatment led to significantly increased mRNA and protein levels of the cytokines IL-1𝛼, IL-1𝛽, and IL-6 in THP-1 monocytes infected with *Legionella* or treated with Pam3CSK4 compared to monocytes not treated with GM-CSF (Figs 1A to 1B). Notably, GM-CSF was insufficient to induce cytokine expression in uninfected cells or cells not stimulated with TLR2 ligand, indicating that a bacterial signal is required for GM-CSF to enhance cytokine production.

**Figure 1.**
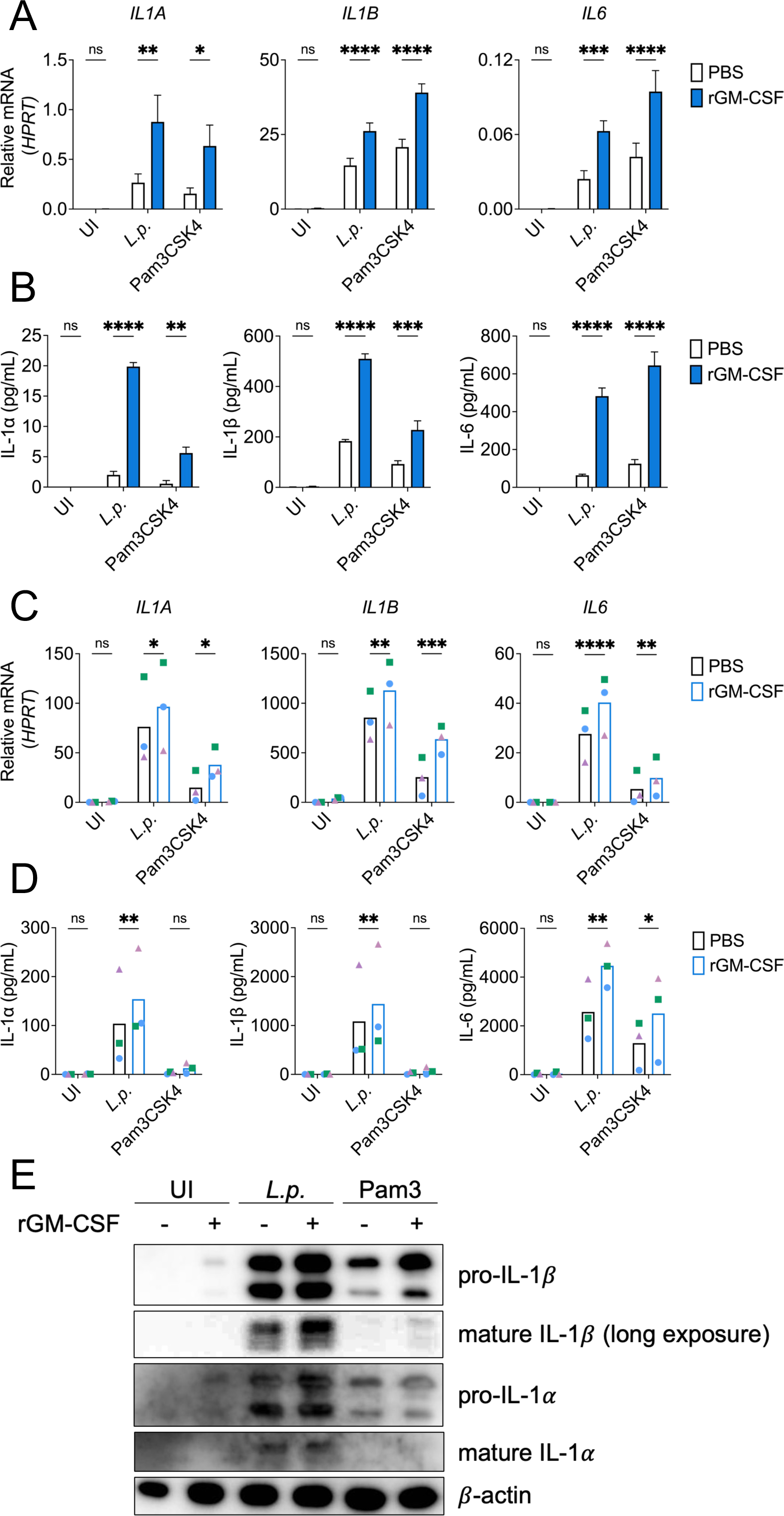
GM-CSF enhances inflammatory responses in human monocytes during *Legionella* infection. (A to B) THP-1 human monocytes or (C to E) primary human monocytes were pre-treated with PBS or rGM-CSF for 30-60min. Cells were then left uninfected, infected with *L.p.* or treated with the TLR2 agonist Pam3CSK4. (A and C) Cells were harvested at 6h after infection to measure *IL1A*, *IL1B*, and *IL6* transcript levels by qPCR, or (B and D) harvested at 24h after infection to measure IL-1𝛼, IL-1𝛽, and IL-6 secretion by ELISA or (E) IL-1𝛼 and IL-1𝛽 protein levels by immunoblot with 𝛽-actin as loading control. Data represent the mean ± SEM of triplicate wells from at least three independent experiments. Data were analyzed by two-way ANOVA with Sidak’s multiple comparisons test; ****, P < 0.0001; ***, P < 0.001; **, P < 0.01; *, P < 0.05; ns, not significant.

We next examined the effect of GM-CSF on cytokine production in primary monocytes isolated from healthy human donors. Consistently, we found elevated IL-1𝛼, IL-1𝛽, and IL-6 transcript and protein levels in the supernatants of *Legionella*-infected or Pam3CSK4-stimulated primary monocytes pretreated with GM-CSF compared to unpretreated monocytes (Figs 1C to 1E). Primary cells treated with both Pam3CSK4 and GM-CSF showed increased intracellular levels of IL-1𝛼 and IL-1𝛽 protein but not released IL-1𝛼 and IL-1𝛽 in supernatants, suggesting that primary human monocytes require an additional signal provided during bacterial infection to release IL-1 cytokines (Fig 1E). Thus, GM-CSF enhances cytokine production in both immortalized and primary human monocytes infected with *Legionella* or stimulated by TLR2 ligands.

We next determined whether GM-CSF enhances the expression of additional cytokine genes (Figure S1). We measured *IL12A* and *IL10* mRNA levels in THP-1 cells treated with or without GM-CSF prior to infection with *Legionella* or stimulation with Pam3CSK4. Although *IL12A* transcript levels were generally low for all conditions, we observed that GM-CSF slightly increased mRNA levels in uninfected cells, but not in *Legionella*-infected or Pam3CSK4-treated cells. GM-CSF pretreatment did not alter *IL10* transcript levels in uninfected or *Legionella*-infected cells, and it decreased *IL10* mRNA levels in cells stimulated with Pam3CSK4.

Collectively, these data indicate that GM-CSF does not universally increase the expression of all cytokine genes and that, similar to what we previously observed in murine monocytes, GM-CSF robustly augments the expression of select inflammatory cytokines in human monocytes infected with *Legionella* or stimulated with TLR2 ligands.

### *Legionella*-induced NF-κB signaling is required for GM-CSF to enhance pro-inflammatory cytokine responses in human monocytes

GM-CSF alone is insufficient to induce inflammatory cytokine expression in uninfected human monocytes (Figure 1). This finding indicates that an additional infection-derived signal is required by monocytes to initiate the expression of cytokines, which is then further enhanced by GM-CSF. Given that GM-CSF promotes cytokine production in monocytes stimulated through TLR2 alone (Figure 1), we hypothesized that initial TLR signaling was required for GM-CSF to augment cytokine expression during *Legionella* infection, as *Legionella* is known to activate TLR signaling (43–50). A major transcription factor that orchestrates cytokine expression downstream of TLR activation and is activated during *Legionella* infection is NF-κB (51–56).

NF-κB signaling is regulated by the multi-subunit IκB kinase (IKK) complex, which phosphorylates IκBα, resulting in its dissociation from NF-κB and its subsequent degradation to enable NF-κB’s translocation into the nucleus where it induces the expression of target genes (51).

Given that GM-CSF’s ability to enhance cytokine production in infected primary human monocytes was largely recapitulated in THP-1 cells, we decided to leverage THP-1 cells as a more tractable system for investigating the role of TLR-dependent NF-κB signaling in this response. To assess activation of NF-κB signaling, we examined IκBα phosphorylation in uninfected or *Legionella*-infected THP-1 cells pretreated with vehicle control or GM-CSF. Although GM-CSF receptor-dependent signaling can lead to NF-κB signaling (57–59), we did not detect IκBα phosphorylation in GM-CSF-treated uninfected cells (Fig 2A). Following *Legionella* infection, we observed a robust increase in IκBα phosphorylation, with no further increase in IκBα phosphorylation with GM-CSF treatment (Fig 2A). These data indicate that *Legionella* infection robustly activates NF-κB signaling in THP-1 cells and that GM-CSF does not further increase NF-κB activation.

**Figure 2.**
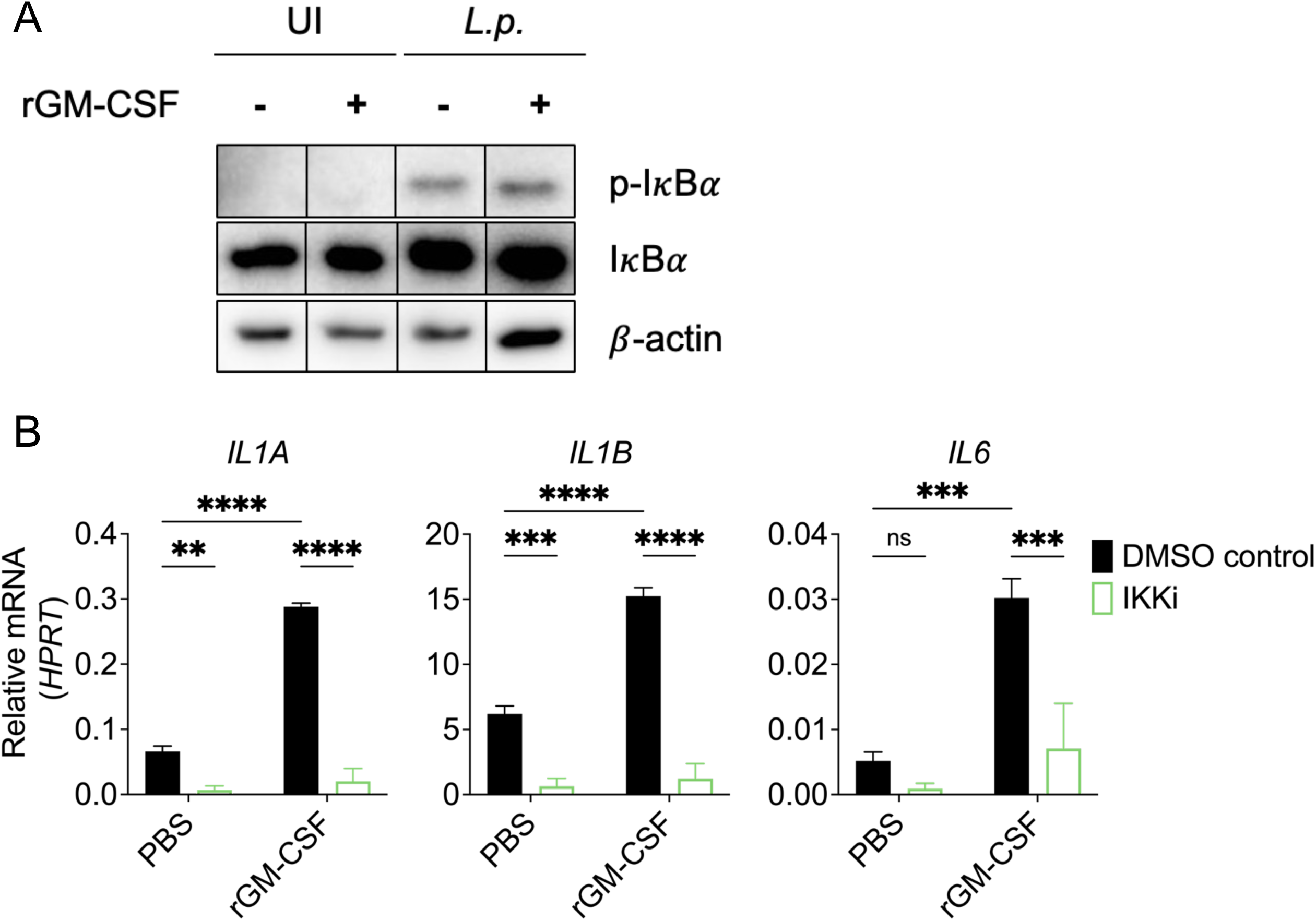
*Legionella*-induced NF-κB signaling is required for GM-CSF-enhanced pro-inflammatory cytokine responses in human monocytes. (A) THP-1 human monocytes were pre-treated with PBS or rGM-CSF for 1hr. Cells were harvested at 6h after infection to perform immunoblot analysis for phospho-I𝜅B𝛼, total I𝜅B𝛼, or 𝛽-actin as loading control. Lanes from one membrane have been cropped and moved to depict the appropriate conditions. No changes were made to the original image during the editing. (B) THP-1 monocytes were pre-treated with vehicle control or the IKK inhibitor BMS-345541 for 1h. Cells were then treated with PBS or rGM-CSF for 1hr followed by *L.p.* infection. Cells were harvested at 6h after infection to measure *IL1A*, *IL1B*, and *IL6* transcript levels by qPCR. Data represent the mean ± SEM of triplicate wells from at least three independent experiments. Data were analyzed by two-way ANOVA with Sidak’s multiple comparisons test; ****, P < 0.0001; ***, P < 0.001; **, P < 0.01; ns, not significant.

To test whether NF-κB signaling is required for GM-CSF to enhance cytokine expression during infection, we treated THP-1 human monocytes with the IKK inhibitor BMS-345541 prior to GM-CSF stimulation and *Legionella* infection. We observed nearly complete abrogation of IκBα phosphorylation and cytokine expression in *Legionella-*infected cells treated with the IKK inhibitor compared to vehicle control (Fig S2 and 2B). Furthermore, GM-CSF failed to increase cytokine expression in *Legionella*-infected cells treated with the IKK inhibitor (Fig 2B).

Altogether, these data indicate that *Legionella*-induced NF-κB signaling is a prerequisite signal for GM-CSF to enhance cytokine expression in human monocytes.

### GM-CSF-dependent JAK2-STAT5 signaling enhances cytokine expression during Legionella infection

We next sought to determine the signaling pathways engaged by GM-CSF to augment inflammatory cytokine expression in monocytes during *Legionella* infection. GM-CSF receptor signaling canonically activates JAK2, which phosphorylates the transcription factor STAT5 and enables its translocation to the nucleus for the induction of target genes (6, 36, 38). We first asked whether there was activation of this pathway by assessing STAT5 phosphorylation in THP-1 monocytes pretreated with GM-CSF or vehicle control prior to *Legionella* or mock infection. We observed STAT5 phosphorylation only when GM-CSF was present (Fig 3A). These data indicate that *Legionella* infection alone does not trigger STAT5 phosphorylation, and that STAT5 signaling is uniquely engaged by GM-CSF in human monocytes.

**Figure 3.**
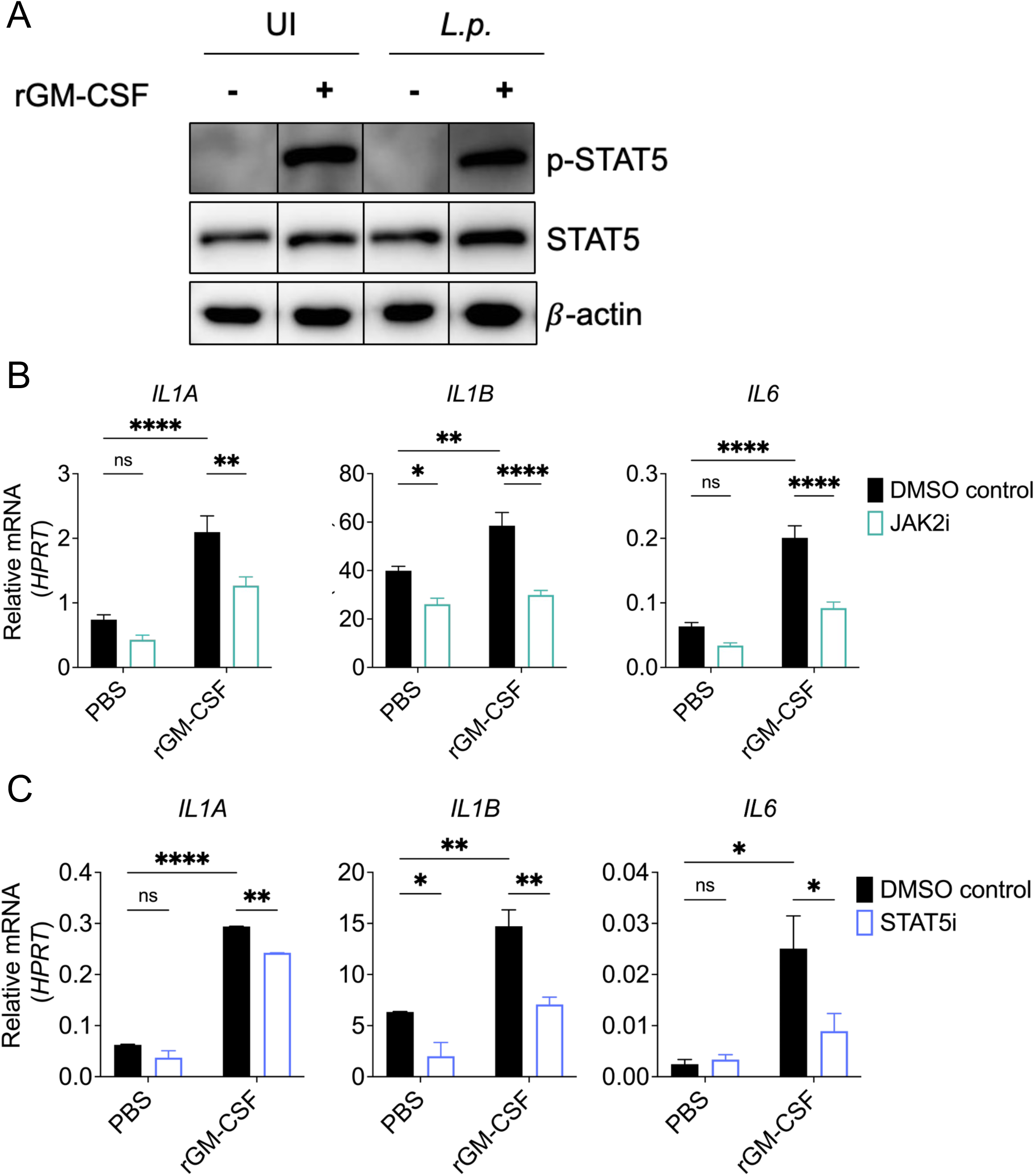
GM-CSF-dependent JAK2-STAT5 signaling is required to enhance inflammatory cytokine expression during *Legionella* infection. (A) THP-1 human monocytes were pre-treated with PBS or rGM-CSF for 1hr. Cells were harvested at 6h after infection to perform immunoblot analysis for phospho-STAT5, total STAT5, or 𝛽-actin as loading control. Lanes from one membrane have been cropped and moved to depict the appropriate conditions. No changes were made to the original image during the editing. (B to C) THP-1 monocytes were pre-treated with vehicle control, (B) the JAK2 inhibitor NVP-BSK805, or (C) the STAT5 inhibitor SH 4-54 for 1h. Cells were then treated with PBS or rGM-CSF for 30-60min followed by *L.p.* infection. Cells were harvested at 6h after infection to measure *IL1A*, *IL1B*, and *IL6* transcript levels by qPCR. Data represent the mean ± SEM of triplicate wells from at least two (C) or three (B) independent experiments. Data were analyzed by two-way ANOVA with Sidak’s multiple comparisons test; ****, P < 0.0001; ***, P < 0.001; **, P < 0.01; *, P < 0.05; ns, not significant.

To test whether JAK2-STAT5 signaling is required for GM-CSF-enhanced cytokine production, we treated THP-1 human monocytes with the JAK2 inhibitor NVP-BSK805 or STAT5 inhibitor SH 4-54 prior to stimulation with GM-CSF and *Legionella* infection. As expected, STAT5 phosphorylation was substantially decreased upon inhibition of JAK2 or STAT5 in *Legionella*-infected THP-1 monocytes treated with GM-CSF compared to vehicle control (Fig S3). We observed a significant decrease in the GM-CSF-dependent upregulation of *IL1A*, *IL1B* and *IL6* in cells treated with the JAK2 or STAT5 inhibitors compared to vehicle control-treated cells, indicating that JAK2-STAT5 signaling is required for GM-CSF to increase expression of these cytokine genes (Fig 3B and 3C). Altogether these data indicate that GM-CSF engages JAK2-STAT5 signaling to enhance inflammatory cytokine responses in human monocytes during *Legionella* infection.

### PI-3K/Akt/mTORC1 signaling is required for GM-CSF-enhanced cytokine expression in *Legionella*-infected human monocytes

GM-CSF can also activate the PI-3K/Akt/mTORC1 pathway (38, 42, 57, 60, 61). Therefore, we determined the contribution of this signaling cascade to GM-CSF-enhanced cytokine responses in infected human monocytes. PI-3Ks are a group of plasma membrane-associated lipid kinases that activate several downstream targets, such as Akt. Akt then promotes activation of mTORC1, which regulates many cellular processes, including the immune response (62–64). We first assessed Akt activation by examining its phosphorylation. Uninfected cells exhibited constitutive Akt phosphorylation, and there was not any detectable increase in *Legionella*- or GM-CSF-dependent Akt activation (Fig S4A). We decided to still examine the contribution of the PI-3K/Akt/mTORC1 pathway in GM-CSF-enhanced cytokine expression of *Legionella*-infected monocytes by assessing its requirement (65). We pretreated cells with the PI-3K inhibitor Ly294002, Akt inhibitor MK-2206, or mTORC1 inhibitor rapamycin and then stimulated with GM-CSF, followed by infection with *Legionella*. Akt phosphorylation was substantially decreased in human monocytes treated with the PI-3K or Akt inhibitor, indicating that these inhibitors were effective (Fig S4B). As expected, Akt phosphorylation was unaffected by mTORC1 inhibition, given that mTORC1 is downstream of Akt (Fig S4B) (62–64). There was a trend toward decreased cytokine mRNA levels, with few statistically significant differences, in PI-3K, Akt, or mTORC1 inhibitor-treated cells infected with *Legionella* in the absence of GM-CSF stimulation (Figs 4A to 4C). We generally observed a significant decrease in cytokine gene expression in GM-CSF-stimulated, *Legionella*-infected monocytes treated with each inhibitor when compared to vehicle control-treated cells (Figs 4A to 4C), although the effect on *IL1B* expression was not significant in cells treated with the PI-3K inhibitor. Given our observation that *IL1B* expression is Akt-dependent, these data suggest that PI-3K may not be the only mechanism supporting Akt-dependent *IL1B* expression. Taken together, these data indicate that PI-3K/Akt/mTORC1 signaling has a minor contribution to *Legionella*-induced cytokine expression but is critical for GM-CSF-dependent induction of cytokine genes in *Legionella*-infected human monocytes.

**Figure 4.**
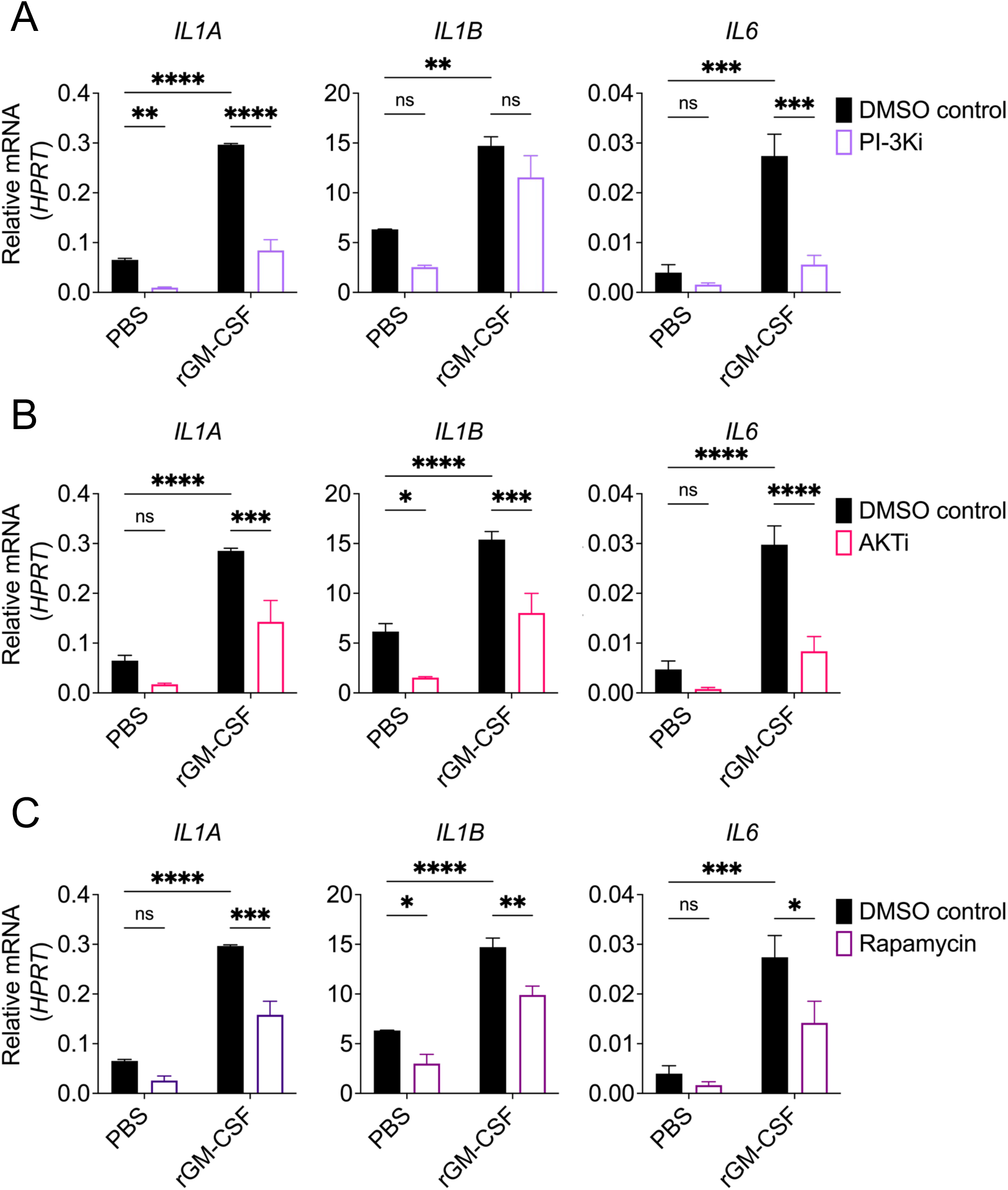
PI-3K/Akt/mTORC1 signaling is required for GM-CSF-dependent pro-inflammatory cytokine expression in *Legionella*-infected human monocytes. (A to C) THP-1 monocytes were pre-treated with vehicle control, (A) PI3-K inhibitor Ly294002, (B) Akt inhibitor MK-2206 or (C) mTORC1 inhibitor Rapamycin for 1h. Cells were then treated with PBS or rGM-CSF for 1hr followed by *L.p.* infection. Cells were harvested at 6h after infection to measure *IL1A*, *IL1B*, and *IL6* transcript levels by qPCR. Data represent the mean ± SEM of triplicate wells from at least three independent experiments. Data were analyzed by two-way ANOVA with Sidak’s multiple comparisons test; ****, P < 0.0001; ***, P < 0.001; **, P < 0.01; *, P < 0.05; ns, not significant.

### GM-CSF-enhanced cytokine expression in *Legionella*-infected human monocytes requires glycolysis and amino acid metabolism

Previous studies from our lab and others have shown that GM-CSF regulates cellular metabolic pathways, including glycolysis, to promote cytokine responses (36, 58, 65). Additionally, TLR, JAK2-STAT5 and PI-3K/Akt/mTORC1 signaling have an important role in activating metabolic processes, such as glycolysis and amino acid metabolism, that regulate immune cell activation (36, 66–78). Therefore, we tested whether glycolysis and amino acid metabolism are required for GM-CSF-enhanced cytokine responses during *Legionella* infection of human monocytes. To assess the role of glycolysis, we cultured THP-1 human monocytes in media containing galactose as a sole carbon source. Galactose is metabolized via the Leloir pathway before entering glycolysis, which results in a substantial reduction in glycolytic flux that effectively inhibits glycolysis (36, 69, 79, 80). We found that *Legionella*-infected THP-1 cells cultured in galactose-containing media had a significant defect in cytokine gene expression following GM-CSF stimulation compared to infected cells cultured in glucose-containing media (Fig 5A). However, there was still some GM-CSF-dependent enhancement of IL1𝛼 and IL-1𝛽 expression in infected THP-1 cells cultured in galactose-containing media (Fig 5A), indicating that other pathways besides glycolysis also contribute.

**Figure 5.**
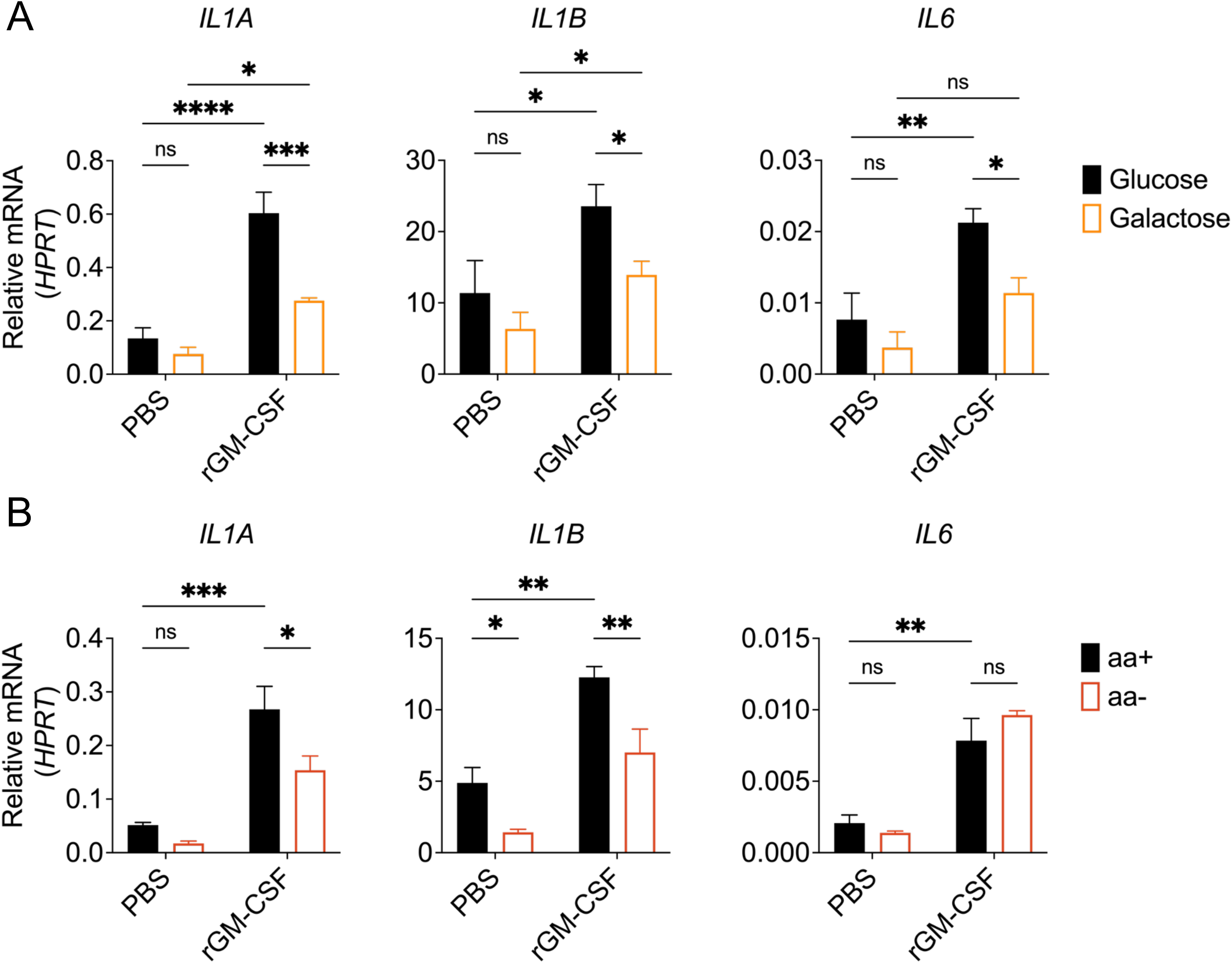
Glycolysis and amino acid metabolism are required for GM-CSF-enhanced pro-inflammatory cytokine responses in human monocytes during *Legionella* infection. (A to B) THP-1 monocytes were replated in (A) media containing glucose or galactose or (B) amino acid sufficient (aa+) or deficient (aa-) media. Cells were then treated with PBS or rGM-CSF for 1hr followed by *L.p.* infection. Cells were harvested at 6h after infection to measure *IL1A*, *IL1B*, and *IL6* transcript levels by qPCR. Data represent the mean ± SEM of triplicate wells from at least three independent experiments. Data were analyzed by two-way ANOVA with Sidak’s multiple comparisons test; ****, P < 0.0001; ***, P < 0.001; **, P < 0.01; *, P < 0.05; ns, not significant.

We previously found that GM-CSF-dependent JAK2-STAT5 signaling in *Legionella-* infected murine monocytes led to increased expression of the genes encoding the glucose transporter GLUT1, hexokinase 2 (*Hk2)*, and phosphofructokinase (*Pfkp*), which are key rate-limiting glycolytic proteins (36). To test whether GM-CSF enhances glycolytic gene expression in *Legionella*-infected THP-1 cells, we measured mRNA levels of *GLUT1*, *HK2*, and *PFKP. Legionella* infection alone did not significantly increase the expression of these glycolytic genes. Additionally, GM-CSF treatment did not increase expression of these genes in uninfected or *Legionella-*infected THP-1 cells (Fig S5), indicating that GM-CSF does not significantly induce transcriptional changes in these glycolytic genes in THP-1 cells (81).

Given that glycolysis does not account for the totality of GM-CSF-enhanced IL-1𝛼 and IL-1𝛽 expression and given our inhibitor data suggesting a role for mTORC1, we sought to interrogate a role for amino acid metabolism by culturing THP-1 monocytes in amino acid-sufficient (aa+) or -deficient (aa-) media. We observed that monocytes cultured in aa-media had reduced *IL1A* and *IL1B* expression after treatment with GM-CSF and *Legionella* infection (Fig 5B). Interestingly, *IL6* transcript levels were similar in cells cultured in aa- or aa+ media, indicating that GM-CSF requires amino acid metabolism to regulate expression of only a subset of its target genes (Fig 5B). Collectively, these data indicate that both glycolysis and amino acid metabolism are required for GM-CSF-dependent enhancement of cytokine expression in human monocytes during infection.

## Discussion

In this study, we investigated the effects of GM-CSF on proinflammatory cytokine production in human monocytes during *Legionella* infection. First, we found that GM-CSF-mediated enhancement of cytokine production requires a pathogen-derived signal that initiates NF-κB-dependent cytokine expression. We also show that JAK2/STAT5 signaling is required for GM-CSF to enhance cytokine expression during infection. Furthermore, we discovered that PI-3K/Akt/mTORC1 signaling is important for GM-CSF-dependent gene induction in human monocytes infected with *Legionella*. Finally, we show that glycolysis and amino acid metabolism are required for GM-CSF to enhance cytokine responses in human monocytes during *Legionella* infection. In summary, these findings show that multiple signaling and metabolic pathways regulate GM-CSF-mediated enhancement of cytokine responses in *Legionella-*infected human monocytes (Figure 6).

**Figure 6.**
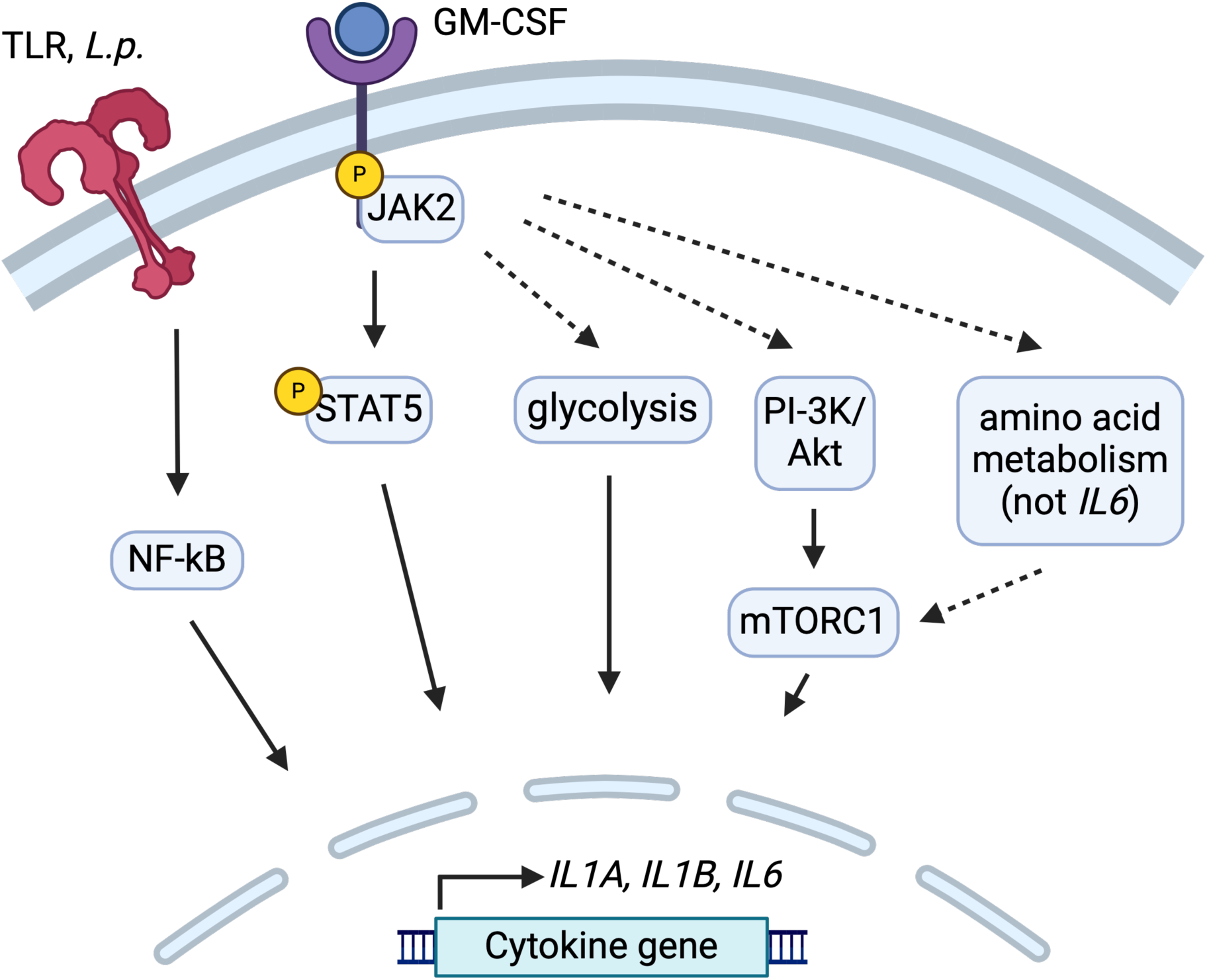
Multistep regulation of GM-CSF-enhanced cytokine gene expression in *Legionella*-infected human monocytes. Graphical model depicting the findings of this study. GM-CSF-enhanced cytokine production requires an initial bacterial PAMP that activates NF-𝜅B signaling in *Legionella*-infected human monocytes. Cytokine gene expression is then enhanced by GM-CSF-dependent JAK2/STAT5 signaling. PI-3K/Akt/mTORC1 signaling, glycolysis, and amino acid metabolism are required for upregulation of cytokine responses by GM-CSF. Dashed lines represent links that need to be further dissected in future studies.

Previous studies demonstrated that GM-CSF enhances pro-inflammatory responses in monocytes in the context of LPS priming (38, 40, 41, 82). However, numerous pathogens have developed strategies to suppress the immune response, underscoring the importance of understanding the role of GM-CSF during active infection. In this study, we employed the bacterial pathogen *Legionella pneumophila*, which uses a type IV secretion system (T4SS) to inject over 300 effectors into the host cell cytosol to manipulate eukaryotic processes, including host signaling and translation (31, 83–87). Despite *Legionella*’s ability to block host protein synthesis, we observed that GM-CSF enhances cytokine mRNA and protein levels in infected human monocytes (Figure 1). Interestingly, GM-CSF is highly induced in murine macrophages in response to *Legionella* blockade of host protein synthesis (31), and IL-1 released by infected macrophages can induce GM-CSF production by alveolar epithelial cells during *Legionella* infection (36). Perhaps GM-CSF is employed by the host to allow immune cells to mount more effective cytokine responses to immunoevasive pathogens.

We found that GM-CSF-mediated enhancement of cytokine production in human monocytes depends on JAK2-STAT5 and PI-3K/Akt/mTORC1 signaling. Additionally, glycolysis and amino acid metabolism are required for GM-CSF to enhance cytokine expression. Whether there is crosstalk between these pathways remains to be carefully dissected. For example, previous research from our lab has shown that JAK2-STAT5 signaling induces the expression of genes encoding the key rate-limiting glycolytic enzymes hexokinase 2 (HK2) and phosphofructokinase (PFKP) in murine monocytes (36). However, we did not observe the upregulation of these genes (Fig S5), suggesting that any potential crosstalk between JAK2-STAT5 signaling and glycolysis occurs through a different mechanism in human monocytes.

Additionally, STAT5 has been shown to regulate amino acid metabolism in T cells by upregulating enzyme and transporter genes (70). STAT5 may also play a role in coordinating amino acid metabolism in GM-CSF-stimulated human monocytes. Another potential link between these pathways is through Akt-mediated activation of glycolysis. Akt can induce glycolysis by upregulating glucose transporter 1 (GLUT1) and lactate dehydrogenase A (LDHA) as well as by translocating HK2 to the mitochondrial outer membrane (78, 88). Furthermore, our findings indicate that mTORC1 is required for GM-CSF-enhanced cytokine expression. mTORC1 can be activated by Akt or amino acid metabolism (89). We showed that GM-CSF-dependent *IL1A*, *IL1B*, and *IL6* expression is Akt- and mTORC1-dependent (Fig 4). In contrast, amino acid metabolism is critical to promote *IL1A* and *IL1B* induction, but not *IL6* (Fig 5B). One potential explanation for this is that depending on the signal mTORC1 might have a different functional outcome. These potential links suggest an intricate interplay between JAK2-STAT5, PI-3K/Akt/mTORC1, glycolysis, and amino acid metabolism pathways in regulating pro-inflammatory cytokine production in human monocytes upon GM-CSF stimulation. Further investigation is needed to unravel the crosstalk between these pathways and how they regulate cytokine expression.

Taken together, our findings demonstrate that human monocytes integrate bacterial and GM-CSF-derived signals to activate multiple signaling pathways, leading to enhanced pro-inflammatory cytokine production. We found that *Legionella*-induced NF-κB activation is a requisite signal for GM-CSF to enhance immune gene expression. Additionally, GM-CSF acts through JAK2-STAT5 signaling to increase cytokine mRNA levels. Furthermore, PI- 3K/Akt/mTORC1 signaling is critical for GM-CSF to augment pro-inflammatory gene transcripts. We also found a requirement for glycolysis and amino acid metabolism in GM-CSF-driven cytokine responses during infection. Collectively, these findings provide a foundation for further understanding how GM-CSF modulates pro-inflammatory responses in human monocytes during infection as well as its role in other inflammatory processes.

## Materials and Methods

### Ethics Statement

All studies on primary human monocytes were performed in compliance with the requirements of the US Department of Health and Human Services and the principles expressed in the Declaration of Helsinki. Samples obtained from the University of Pennsylvania Human Immunology Core are considered to be a secondary use of deidentified human specimens and are exempt via Title 55 Part 46, Subpart A of 46.101 (b) of the Code of Federal Regulations.

### Cell Culture

THP-1 cells (TIB-202; American Type Culture Collection) were maintained in RPMI supplemented with 10% (vol/vol) heat-inactivated FBS, 0.05 mM β-mercaptoethanol, 100 IU/mL penicillin, and 100 μg/mL streptomycin at 37 °C in a 5% CO2 humidified incubator. One day before infection, cells were replated in media lacking antibiotics at a concentration of 2.0 × 10^5^ cells per well in a 48-well plate.

Primary human monocytes from deidentified healthy human donors were obtained from the University of Pennsylvania Human Immunology Core. Monocytes were cultured in RPMI supplemented with 10% (vol/vol) heat-inactivated FBS, 2 mM L-glutamine, 100 IU/mL penicillin and 100 μg/mL streptomycin. One day before infection, cells were replated in media lacking antibiotics at a concentration of 2.0 × 10^5^ cells per well in a 48-well plate or 4×10^5^ cells per well in a 96-well plate.

For TLR2 stimulation, cells were treated with 10ng/mL Pam3CSK4 (Invivogen tlrl-pms), For infections with *L. pneumophila*, bacteria were resuspended in PBS and added to the cells at a multiplicity of infection (MOI) of 5. The plate was then centrifuged at 1200rpm for 5 min and placed into a humidified tissue culture incubator set at 37°C, 5% CO2. Cells were infected for 6 h for analysis by qPCR and 24 h for analysis by ELISA or Immunoblot.

For GM-CSF stimulation, cells were treated with 10ng/mL of human GM-CSF (Biolegend 572902) 1hr prior to Pam3CSK4 stimulation or *Legionella* infection.

For inhibitor treatments, cells were pre-treated 1hr before GM-CSF stimulation with either 250nM IKK inhibitor (BMS-345541; Selleck Chemicals S8044), 2uM JAK2 inhibitor (NVP-BSK805; Selleck Chemicals S2686), 5uM STAT5 inhibitor (SH 4-54; Selleck Chemicals S7337), 10uM PI-3K inhibitor (Ly294002; Selleck Chemicals S1105), 5uM Akt inhibitor (MK- 2206; Selleck Chemicals S1078), or 100nM Rapamycin (AY-22989; Selleck Chemicals S1039).

For experiments to probe glycolysis, THP-1 monocytes were incubated in media containing glucose (10 mM) or galactose (10 mM) and were then infected with *L.p.* (MOI = 5) for 6 hr. For experiments to probe amino acid metabolism, THP-1 monocytes were incubated in amino acid-free RPMI 1640 (MyBioSource MBS652918) that was or was not reconstituted with each individual amino acid at concentrations equivalent to those contained in standard RPMI 1640 followed by infection with *L.p.* (MOI = 5) for 6 hr.

### Bacterial Cultures

*Legionella pneumophila* serogroup 1 strains were used in all experiments. Primary human monocytes or THP-1 cells were infected with a *Legionella pneumophila* Lp02-derived (thymidine auxotroph) *ΔflaA* mutant (90, 91). *L pneumophila* was cultured on charcoal yeast extract (CYE) agar plates for 48 h at 37°C prior to infection.

### Reverse transcription-quantitative PCR (qRT-PCR) analysis

Cells were lysed, and RNA was isolated using the RNeasy Plus Kit (Qiagen). Synthesis of the first strand cDNA was performed using SuperScript II reverse transcriptase and oligo (dT) primer (Invitrogen). qPCR was performed with the CFX96 real-time system (Bio-Rad) using the SsoFast EvaGreen Supermix with the Low ROX kit (Bio-Rad). The following primers were used. *IL1A* forward: TGGTAGTAGCAACCAACGGGA, *IL1A* reverse:

ACTTTGATTGAGGGCGTCATTC, *IL1B* forward: AGCTACGAATCTCCGACCAC, *IL1B* reverse: CGTTATCCCATGTGTCGAAGAA, *IL6* forward: ACTCACCTCTTCAGAACGAATTG, *IL6* reverse: CCATCTTTGGAAGGTTCAGGTTG,*HPRT* forward: CCTGGCGTCGTGATTAGTGAT, *HPRT* reverse: AGACGTTCAGTCCTGTCCATAA. The following primers from PrimerBank were used (92–94). The PrimerBank identifications are *GLUT1* (166795298c1), *HK2* (40806188c3), and *PFKP* (334191700c2): *GLUT1* forward: GGCCAAGAGTGT GCTAAAGAA, *GLUT1* reverse: ACAGCGTTGATGCCAGACAG, *HK2* forward: TTGACCAGGAGATTGACATGGG, *HK2* reverse: CAACCGCATCAGGACCTCA, PFKP forward: CGCCTACCTCAACGTGGTG, PFKP reverse: ACCTCCAGAACGAAGGTCCTC.

### Immunoblot Analysis

Infected or treated cells were lysed directly with 1× SDS/PAGE sample buffer. Protein samples were boiled for 5 min, separated by SDS/PAGE, and transferred to PVDF Immobilon-P membranes (Millipore). Samples were then probed with antibodies specific for IL-1𝛼 (R&D Systems MAB200), IL-1β (R&D Systems MAB201), IκBα (Cell Signaling 9242), phospho- IκBα (Cell Signaling 2859), Stat5 (Cell Signaling 94205S), phospho-Stat5 (Cell Signaling 9351S), Akt (Cell Signaling 9272), and phospho-Akt (Cell Signaling 9271). As a loading control, all blots were probed with anti–β-actin (Cell Signaling 4967L). Detection was performed with HRP-conjugated anti-mouse IgG (Cell Signaling F00011) or anti-rabbit IgG (Cell Signaling 7074S).

### ELISA

Harvested supernatants from infected cells were assayed using ELISA kits for human IL- 1α (R&D Systems), IL-1β (BD Biosciences), and IL-6 (Biolegend).

### Statistical Analyses

All graphed data and ANOVA analyses were carried out in GraphPad Prism (San Diego, California). ANOVA was followed by multiple comparison with the Holm-Šídák’s post-test. The resulting significance levels are indicated in the figures. All p-values and significance levels are indicated in the figures and figure legends.

## Acknowledgements

We thank members of the Shin and Brodsky laboratories for helpful scientific discussions. We thank Marisa Egan for providing THP-1 monocytes. We thank the Human Immunology Core of the Penn Center for AIDS Research and Abramson Cancer Center for providing purified primary human monocytes. We thank the Penn Summer Undergraduate Internship Program (SUIP) for supporting our summer undergraduate Madison Dresler. This work is supported in part by grants R01AI118861 (S.S.), R01AI123243 (S.S.), R21AI151476 (S.S.), R25HL084665 (M.V.D.), National Science Foundation Graduate Fellowship DGE- 1845298 (V.R.V.M), American Heart Association Predoctoral Fellowship 23POST1011760 (M.D.H.), and a Burroughs-Welcome Fund Investigators in the Pathogenesis of Infectious Diseases Award (S.S.).

## Supplemental Figures

**Figure S1** GM-CSF does not upregulate expression of any inflammatory cytokine genes. THP-1 monocytes were pre-treated with PBS or rGM-CSF for 30-60min. Cells were then left uninfected, infected with *L.p.* or treated with the TLR2 agonist Pam3CSK4. Cells were harvested at 6h after infection to measure *IL12A* and *IL10* transcript levels by qPCR. Data represent the mean ± SEM of triplicate wells from at least three independent experiments. Data were analyzed by two-way ANOVA with Sidak’s multiple comparisons test; *, P < 0.05; ns, not significant.

**Figure S2** IKK inhibitor abrogates *Legionella*-induced I𝜅B𝛼 phosphorylation in THP-1 human monocytes. THP-1 monocytes were pre-treated with vehicle control or the IKK inhibitor BMS- 345541 for 1h. Cells were then treated with PBS or rGM-CSF for 1hr followed by *L.p.* infection. Cells were harvested at 6h after infection to perform immunoblot analysis for phospho-I𝜅B𝛼, total I𝜅B𝛼, or 𝛽-actin as loading control. Lanes from one membrane have been cropped and moved to depict the appropriate conditions. No changes were made to the original image during the editing.

**Figure S3** STAT5 and JAK2 inhibitors abrogate GM-CSF-dependent STAT5 phosphorylation in THP-1 human monocytes infected with *Legionella*. THP-1 monocytes were pre-treated with vehicle control, the JAK2 inhibitor NVP-BSK805, or the STAT5 inhibitor SH 4-54 for 1h. Cells were then treated with PBS or rGM-CSF for 30-60min followed by *L.p.* infection. Cells were harvested at 6h after infection to perform immunoblot analysis for phospho-STAT5, total STAT5, or 𝛽-actin as loading control.

**Figure S4** THP-1 human monocytes exhibit basal Akt phosphorylation that is downregulated by PI-3K and AKT inhibitors. (A) THP-1 human monocytes were pre-treated with PBS or rGM-CSF for 1hr. Cells were harvested at 6h after infection to perform immunoblot analysis for phospho-Akt, total Akt, or 𝛽-actin as loading control. Lanes from one membrane have been cropped and moved to depict the appropriate conditions. No changes were made to the original image during the editing. (B) THP-1 monocytes were pre-treated with vehicle control, PI3-K inhibitor Ly294002, Akt inhibitor MK-2206 or mTORC1 inhibitor Rapamycin for 1h. Cells were then treated with PBS or rGM-CSF for 1hr followed by *L.p.* infection. Cells were harvested at 6h after infection to perform immunoblot analysis for phospho-Akt, total Akt, or 𝛽-actin as loading control. Lanes from one membrane have been cropped and moved to depict the appropriate conditions. No changes were made to the original image during the editing.

**Figure S5** GM-CSF does not significantly upregulate rate-limiting glycolytic genes in *Legionella*-infected human monocytes. THP-1 monocytes were pre-treated with PBS or rGM-CSF for 30-60min. Cells were then left uninfected, infected with *L.p.* or treated with the TLR2 agonist Pam3CSK4. Cells were harvested at 6h after infection to measure *GLUT1, PFKP,* and *HK2* transcript levels by qPCR. Data represent the mean ± SEM of triplicate wells from at least three independent experiments. Data were analyzed by two-way ANOVA with Sidak’s multiple comparisons test; ns, not significant.

