## Supplementary figures and images for "GM-CSF engages multiple signaling pathways to enhance pro-inflammatory cytokine responses in human monocytes during *Legionella* infection"

### Supplemental figures

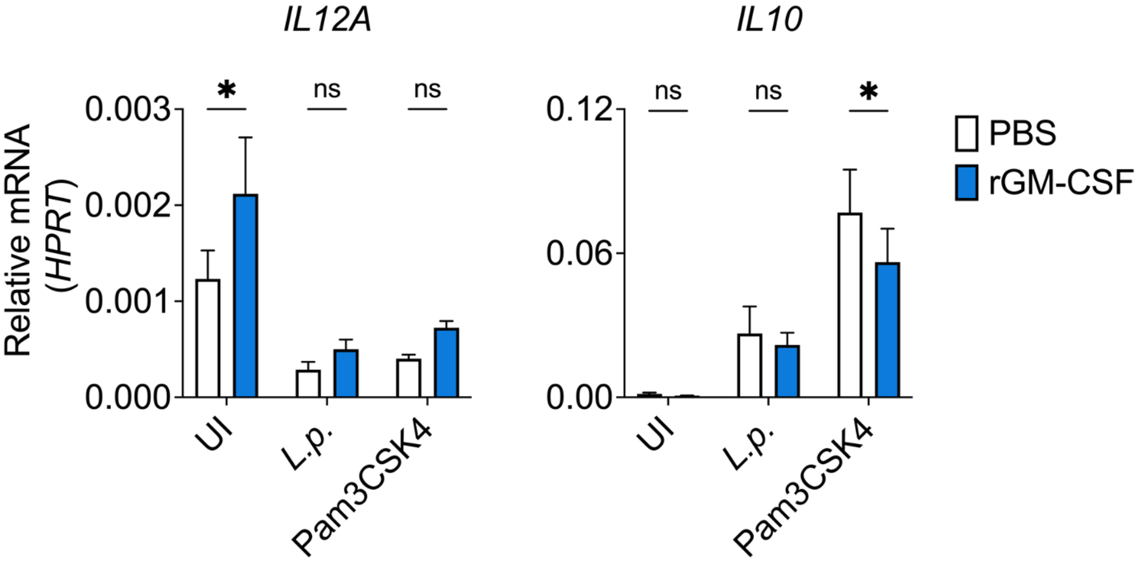

#### Figure S2

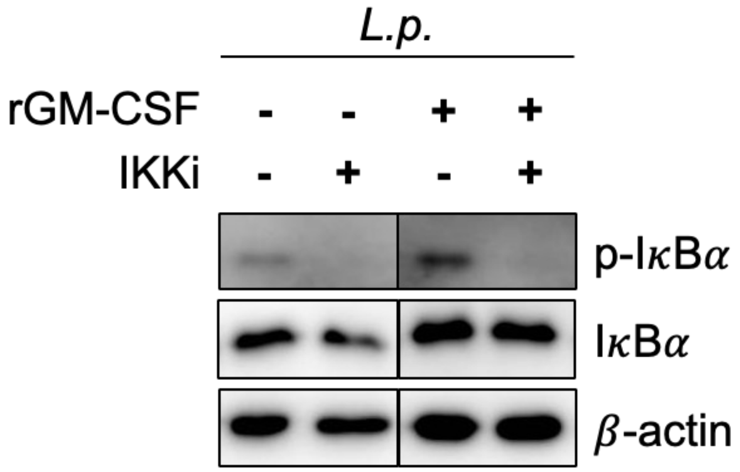

Figure S3

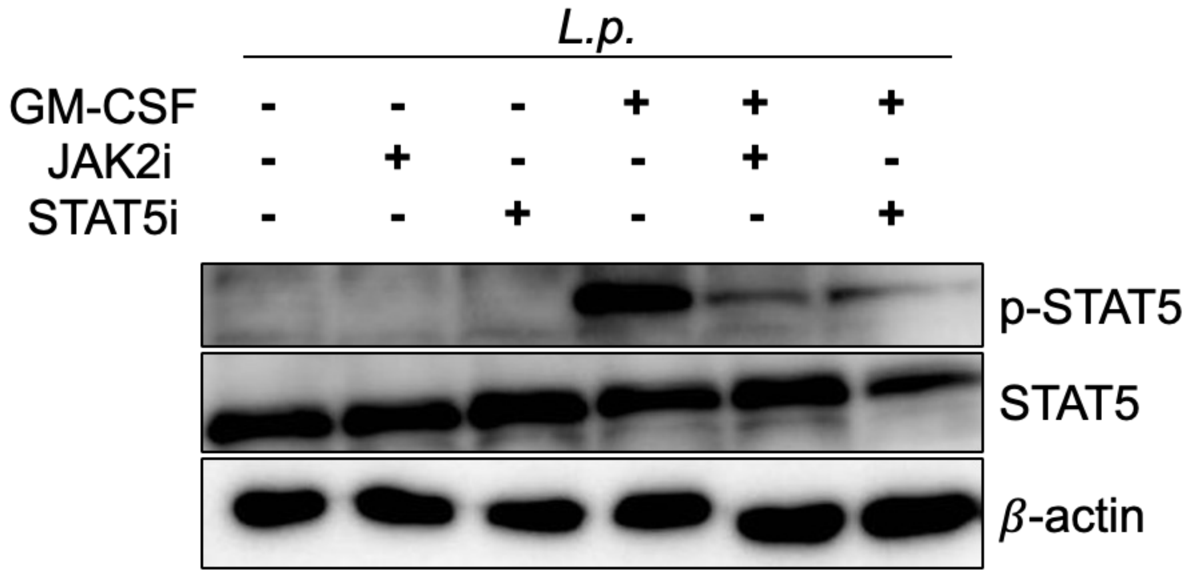

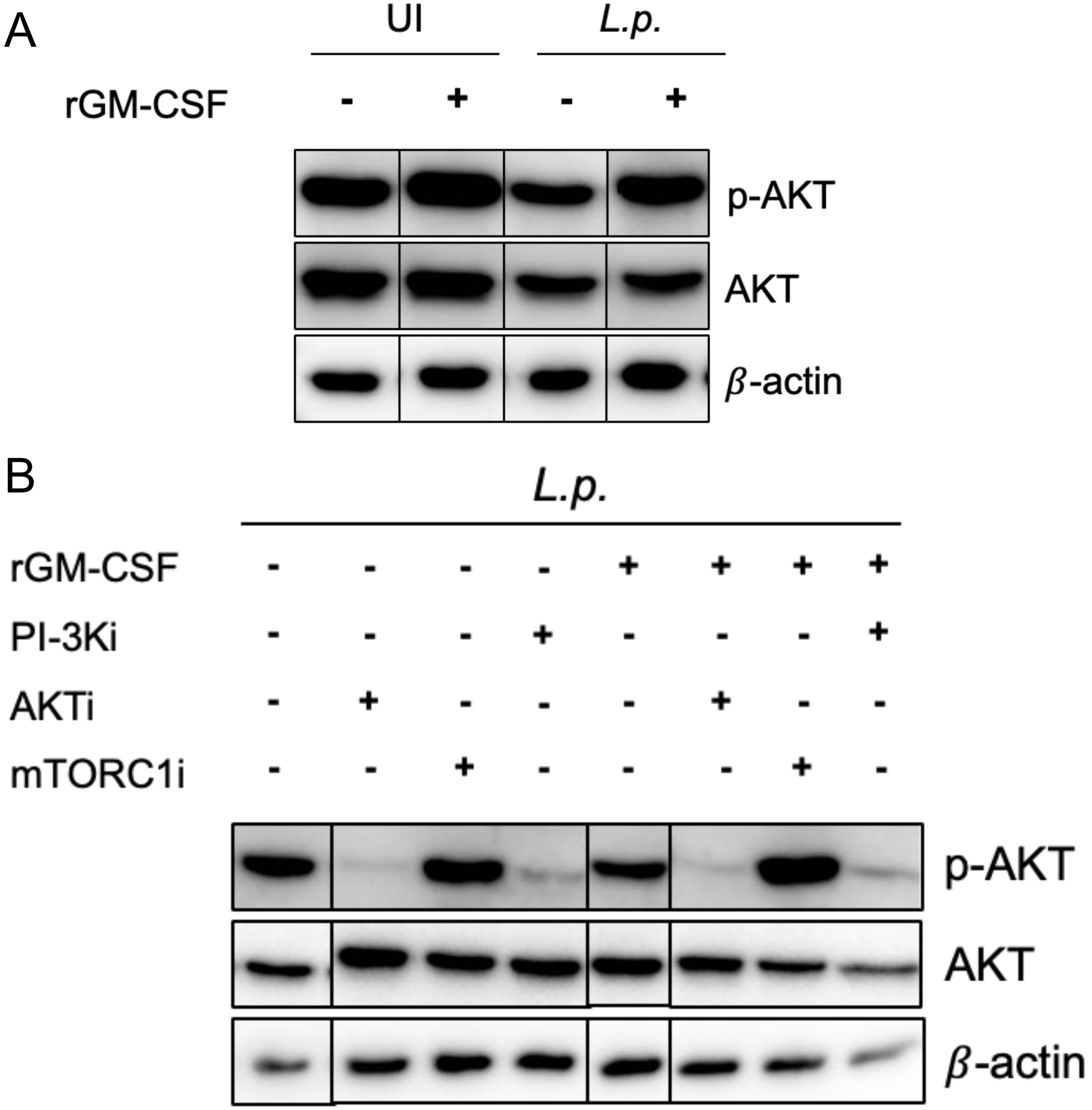

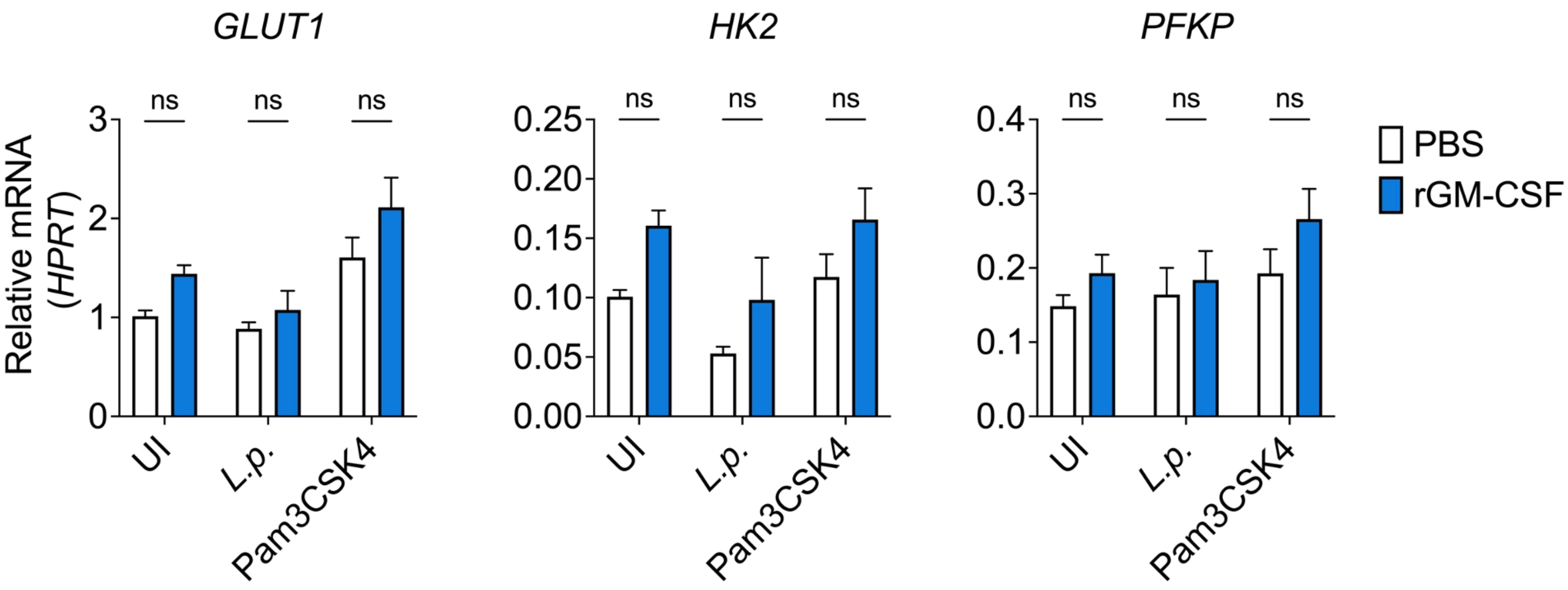
